# Transdiagnostic symptomatology amidst real-world environmental uncertainty: a cross-sectional and cross-lagged panel network analysis

**DOI:** 10.1101/2025.08.25.672142

**Authors:** Friederike Elisabeth Hedley, Claudia Lage, Nimrod Hertz-Palmor, Timothy R. Sandhu, Susanne Schweizer, Rebecca P. Lawson

## Abstract

**Background:** Uncertainty is a potential driver of poor mental health outcomes, and uncertainty is mounting globally across many domains of daily life. However, it remains unclear how anxiety and depression symptoms emerge in response to uncertainty in these real-life contexts. Capitalising on the COVID-19 pandemic as a naturalistic experiment, we investigate how heightened real-world environmental uncertainty interacts with intolerance of uncertainty (IU) to drive mental health symptoms.

**Methods:** We collected self-reported data on symptoms of anxiety, depression and IU from 301 participants for two time points. We performed cross-sectional network analyses to identify influential contributors and established a cross-lagged panel network (CLPN) to investigate longitudinal effects.

**Results:** Anxiety, depression and IU symptoms were higher when real-world uncertainty was higher (*t*2) compared to when it was lower (*t*1). Network analyses revealed the same three symptom clusters at both timepoints, however worry became more central to the network at *t*2. In the CLPN, unlike anxiety and depression symptoms, IU symptoms had the highest autoregressive values, meaning IU at *t*2 was well predicted by IU at *t*1. The item “inhibition of behaviour due to doubt” positively predicted both anxiety and anhedonia.

**Conclusions:** This study highlights the relevance of worry during higher real-world uncertainty and the predictive value of predispositional IU. Our findings further suggest behavioural inhibition to be a potential target to alleviate internalising psychopathology. During ever-increasing fluctuations in global uncertainty, we provide novel insights into the temporal relationships of highly prevalent psychopathology, which might inform support strategies for people with existing mental health vulnerabilities.

## Introduction

Uncertainty is increasingly recognised as a key driver of poor mental health outcomes (Kienzler et al., 2022; Schweizer et al., 2023). Cognitive models of internalising psychopathology, such as anxiety and depressive disorders, highlight the role of uncertainty misestimation and intolerance of uncertainty in both the onset and maintenance of these conditions (Carleton, 2016; Grupe & Nitschke, 2013). Individual differences in how uncertainty is estimated and represented computationally in the brain have been proposed as risk factors for the development of specific mental health conditions (Sandhu et al., 2023), but to fully understand how different psychopathologies emerge, it is important to integrate information across multiple levels of analysis, spanning genetics, neural circuits, behaviour, self-reports, and environmental context (Insel et al., 2010). Here, we focus on a particularly salient but under-examined level: the external environment. We live in a world with pervasive and escalating global uncertainties–including economic instability, political upheaval, armed conflicts, and the climate crisis–that form the backdrop of daily experience for many. However, how such real-world environmental uncertainty contributes to the emergence and maintenance of anxiety and depression, and how it interacts with intolerance of uncertainty, remains poorly understood.

One recent example of heightened global uncertainty was the COVID-19 pandemic, a period marked by a massive surge in the World Uncertainty Index (**Figure 1a**; Ahir et al. (2022)). Fluctuating confinement regulations and evolving epidemiological information contributed to uncertainty across health, occupational, and social domains, arguably affecting mental wellbeing globally (Ahir et al., 2022; Altig et al., 2020; Hertz-Palmor et al., 2023). This period also saw a marked increase in the prevalence of depressive and anxiety symptoms and disorders (Hertz-Palmor et al., 2021; McBride et al., 2020; Santomauro et al., 2021; van der Velden et al., 2021; World Health Organization, 2022) and their contribution to Disability-Adjusted Life Years (DALYs, **Figure 1b**; Global Burden of Disease Collaborative Network). Though the immediate crisis has passed, the pandemic provides a compelling naturalistic experiment illustrating how large-scale environmental uncertainty can fuel the onset and persistence of psychopathology.

**Figure 1.**
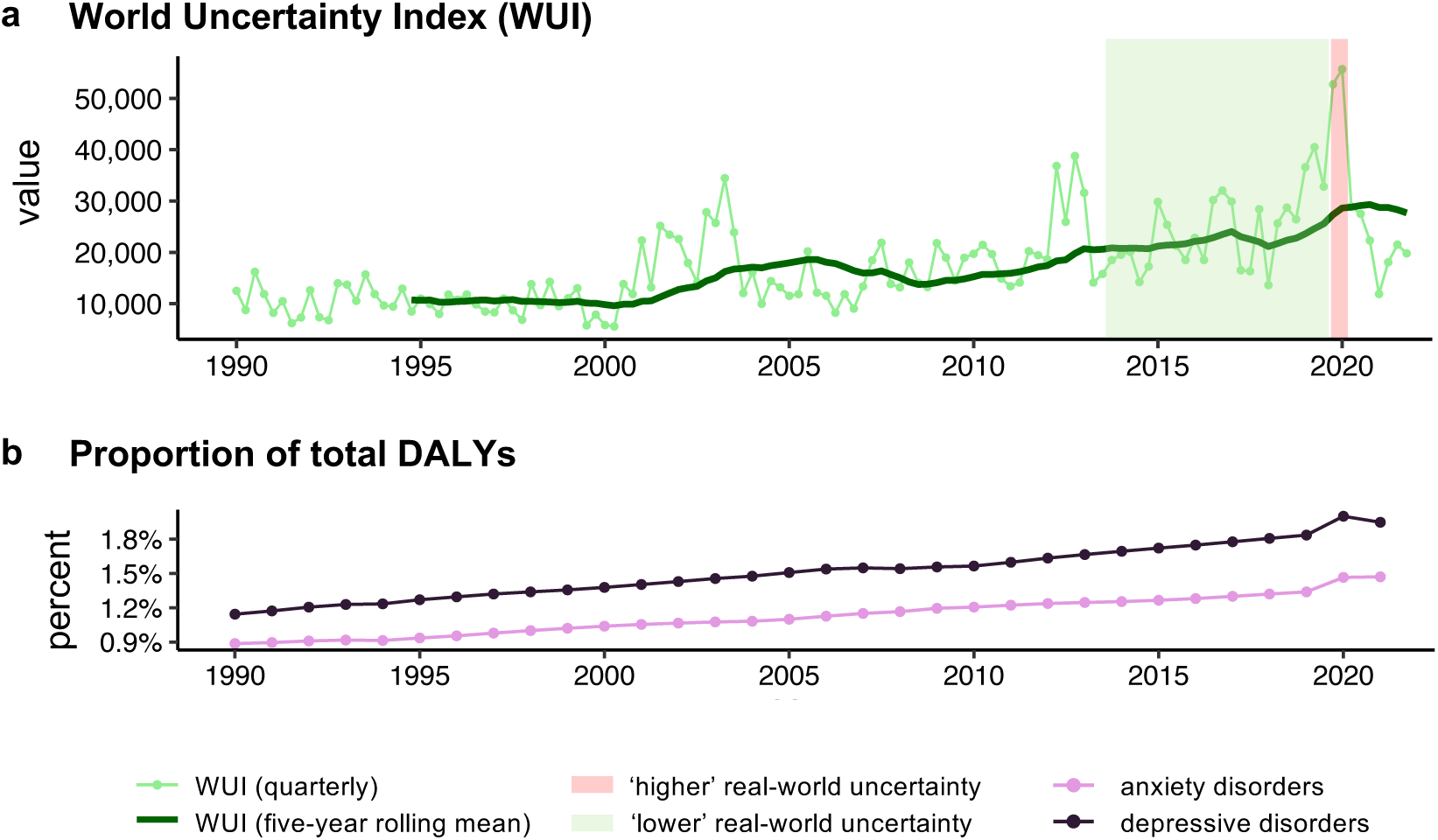
World Uncertainty Index (WUI) and Disability-Adjusted Life Years (DALYs) for anxiety and depressive disorders. **a** World Uncertainty Index, presented in quarterly intervals (light green) and as a five-year rolling mean (dark green), between 1990 and 2021 (Ahir et al., 2022). The index is GDP-weighted for 142 countries. The green and red shaded sections indicate the periods we operationalise as ‘lower’ and ‘higher’ real-world environmental uncertainty, respectively. Note that we chose these labels to juxtapose the two periods, indicating that they are comparative. **b** Proportion of total Disability-Adjusted Life Years (DALYs) for anxiety (pink) and depressive (purple) disorders between 1990 and 2021. Data is sourced from the Institute for Health Metrics and Evaluation (IHME). GBD Compare Data Visualization. Global Burden of Disease Collaborative Network Study 2021. Seattle, WA: IHME, University of Washington, 2023. Available from http://vizhub.healthdata.org/gbd-compare.

Yet not everyone responds to uncertainty in the same way. A key factor shaping individual vulnerability is *intolerance of uncertainty* (IU), a dispositional tendency to react negatively to uncertain situations across emotional, cognitive, and behavioural domains (Carleton, 2016; Dugas et al., 2004). Higher IU is associated with threat overestimation and increased likelihood of maladaptive coping strategies such as rumination, avoidance, or reassurance seeking. As a transdiagnostic construct, IU is strongly linked to anxiety disorders, depressive disorders and obsessive-compulsive disorder (Boswell et al., 2013; Dugas et al., 1998; Hedley et al., 2024; McEvoy et al., 2019; Rosser, 2019), and is increasingly recognised as an important treatment target (Dugas et al., 2022; Robichaud, 2013; Zemestani et al., 2021). IU can be further subdivided into prospective IU, which reflects the discomfort when anticipating uncertain future events and the desire for predictability, and inhibitory IU, which reflects avoidance behaviour or decision-making paralysis in the face of uncertainty (Carleton et al., 2007). Together, IU is a dispositional trait, an important risk factor, and a promising treatment target that is central to research on mental disorders, particularly anxiety and depression.

Traditionally, symptom- and disorder-focused approaches have been used to evaluate the structure of mental disorders (American Psychiatric Association DSM-5 Task Force, 2013; Kessler et al., 2005; World Health Organization, 2019). While these classifications and taxonomies have advanced our understanding of psychopathology, they risk obscuring the complex interplay between symptoms and may fail to capture the dynamic processes that drive comorbidity and clinical outcomes. Network theory offers an alternative perspective on psychopathology, viewing mental disorders as systems of directly interacting symptoms (Borsboom, 2017). In this framework, poor mental health may arise within a dynamical system, for example when strong links between symptoms (e.g., depressed mood and worry) form tightly connected nodes within a cross-sectional network. These relationships–or *edges*–are estimated while controlling for all other symptoms, revealing unique associations. Clustering algorithms can identify symptom communities (e.g., anxious arousal and worry) and cross-cluster connections (e.g., worry and avoidance of surprise), highlighting unique transdiagnostic patterns (Groen et al., 2019; Newman, 2006). A key strength of this approach lies in identifying bridging symptoms and edges that connect distinct clusters, helping to explain comorbidity (Cramer et al., 2010). For example, a recent study showed that during the COVID-19 pandemic, depression-anxiety symptom connections intensified over time among individuals high in IU (Andrews et al., 2023). However, cross-sectional networks cannot fully capture the *temporal* dynamics of symptom change.

Understanding how symptom relationships evolve over time–particularly in response to shifting environmental contexts such as global uncertainty–requires temporal network approaches. Cross-lagged panel network (CLPN) modelling is an emerging method that allows for an analysis of longitudinal effects between symptoms (Wysocki et al., 2022). CLPN models can indicate the direction of potentially causal relationships by examining how nodes (symptom variables) at one timepoint influence symptoms at a subsequent timepoint. In temporal networks, *in-strength* measures a node’s (i.e., symptom) predictability from incoming edges from other nodes at the previous time point, while *out-strength* reflects its influence on other nodes via outgoing edges to the next time point. Prior work has applied CLPN approaches to anxiety and depression symptom networks during COVID-19 quarantine periods, revealing, for example, that “nervousness” exhibited highest predictability (Ramos-Vera et al., 2024). However, it remains unclear how global uncertainty interacts with IU to drive mental health symptoms, a question that requires data reported during periods of lower and higher environmental uncertainty.

In the present study, we investigated how relationships between IU and anxiety and depression symptoms changed in response to real-world environmental uncertainty. Specifically, we first performed cross-sectional network analysis for two timepoints, *lower* and *higher* uncertainty (*t*1 and *t*2, respectively) operationalised as *before* and *during* the COVID-19 pandemic. Our aim was to identify the most influential contributors within networks comprising IU and internalising symptoms, and to compare network structures across time. Second, we used CLPN modelling to examine how symptoms influence one another–and themselves–across the transition from lower to higher real-world environmental uncertainty. By combining cross-sectional and cross-lagged network analyses, our goal was to provide novel insights into the temporal dynamics linking IU to anxiety and mood disorder symptoms, and to identify potential intervention targets for improving mental health outcomes in response to environmental changes.

## Methods

### Participants

Participants completed a survey administered on the Research Electronic Data Capture (REDCap; Harris et al. (2019); Harris et al. (2009)), and anonymised data was collected between April and May 2020. As there no straight forwarded approach for power calculations in psychological networks (Epskamp et al., 2018), we based our sample size on previous network modelling work using IU measures, targeting a minimum of 250 participants (Jach & Smillie, 2019; Ren et al., 2020). After removal of incomplete datasets (*N* = 123), our final sample included 301 participants, predominantly female (77.74%), with ages ranging from 20 to 81 years (*M* = 41.2, *SD* = 14.3), mostly from the United Kingdom (70.76%). Summary statistics for demographic information are presented in **Table 1**. Ethical approval for the study was granted by the Cambridge Psychology Research Ethics Committee (PREC, PRE.2019.110.) Participants’ informed consent was obtained prior to data collection through an online form.

**Table 1.** Demographic characteristics.

| *Characteristics* | *M* | *SD* |
| --- | --- | --- |
| **Age** | 41.2 | 14.3 |
|  | *n* | % |
| **Gender** |  |  |
| Women | 234 | 77.74 |
| Men | 66 | 21.93 |
| Non-binary | 1 | 0.33 |
| **Ethnicity** |  |  |
| English / Welsh / Scottish / Northern Irish / British | 213 | 70.76 |
| Irish | 14 | 4.65 |
| Any other White background | 44 | 14.62 |
| White and Black Caribbean | 3 | 1 |
| White and Asian | 7 | 2.33 |
| Any other Mixed / Multiple ethnic background | 2 | 0.66 |
| Indian | 5 | 1.66 |
| Chinese | 6 | 1.99 |
| Any other Asian background | 2 | 0.66 |
| African | 1 | 0.33 |
| Any other ethnic group | 4 | 1.33 |
| **Context-related measures** |  |  |
| Lockdown initiated | 287 | 95.34 |
| Social distancing recommended | 298 | 99.00 |
| COVID symptoms currently experienced | 3 | 1.00 |

*Note:* Under context-related measures, participants were asked to provide further information regarding the heightened real-world uncertainty context they experienced at the point of study administration (i.e., during the COVID-19 pandemic).

### Measures

Participants completed several questionnaires (see details below) and for each we asked them to indicate their answers for two timepoints. For anxiety and depression measures, we captured their responses for “the last two weeks” (i.e., during the COVID-19 pandemic) and “the two weeks before the Covid-19 outbreak”. For personality trait measures, to anchor participants’ pre-pandemic responses in a common and interpretable way, they were instructed to “indicate how much these statements were characteristic of you before the Covid-19 outbreak”. This kind of retrospective or trait-based self-report is widely used and well-established in clinical and psychological research. For example, instruments such as the Trait subscale of the State-Trait Anxiety Inventory (STAI-T), the Intolerance of Uncertainty Scale (IUS), and the Parental Bonding Instrument (PBI) ask individuals to report either on their typical or characteristic experiences or to recall earlier periods of life. While retrospective reporting may carry some risk of recall bias and mood-related memory confounds (Bower, 1987; Coughlin, 1990; Faul & LaBar, 2023), it remains a valid and commonly accepted approach for capturing individual differences across contexts (American Psychiatric Association DSM-5 Task Force, 2013; Moffitt et al., 2010; Talari & Goyal, 2020). In our analyses, we refer to responses for the period before the pandemic as reflecting *lower* real-world uncertainty (*t*1), and those during the pandemic as reflecting *higher* real-world uncertainty (*t*2).

#### Patient Health Questionnaire

We first report data of the 4-item Patient Health Questionnaire (PHQ-4; Kroenke et al. (2009)), for which responses are recorded on a Likert-scale, ranging from 0 = *not at all* to 3 = *nearly every day*. The PHQ-4 includes two subscales: an anxiety subscale measuring self-reported anxiety/nervousness and worry, and a depression subscale measuring depression/hopelessness and anhedonia. Cut-off scores of ≥3 for the two subscales are suggestive of clinical anxiety and depression, respectively.

#### Intolerance of Uncertainty Scale

We also collected self-reported data from the short-version of the Intolerance of Uncertainty Scale (IUS-12; Carleton et al. (2007)), for which responses are recorded on a Likert-scale, ranging from 1 = *not at all characteristic of me* to 5 = *entirely characteristic of me*. The IUS-12 includes two subscales: a 7-item subscale measuring prospective IU, which reflects the desire for predictability and reduced uncertainty (e.g., “Unforeseen events upset me greatly”), as well as a 5-item subscale measuring inhibitory IU, which captures an individual’s propensity to display avoidance behaviour when faced with uncertainty (e.g., “When it’s time to act, uncertainty paralyses me”). Both measurements and all subscales are shown in **Table 2**.

**Table 2.**
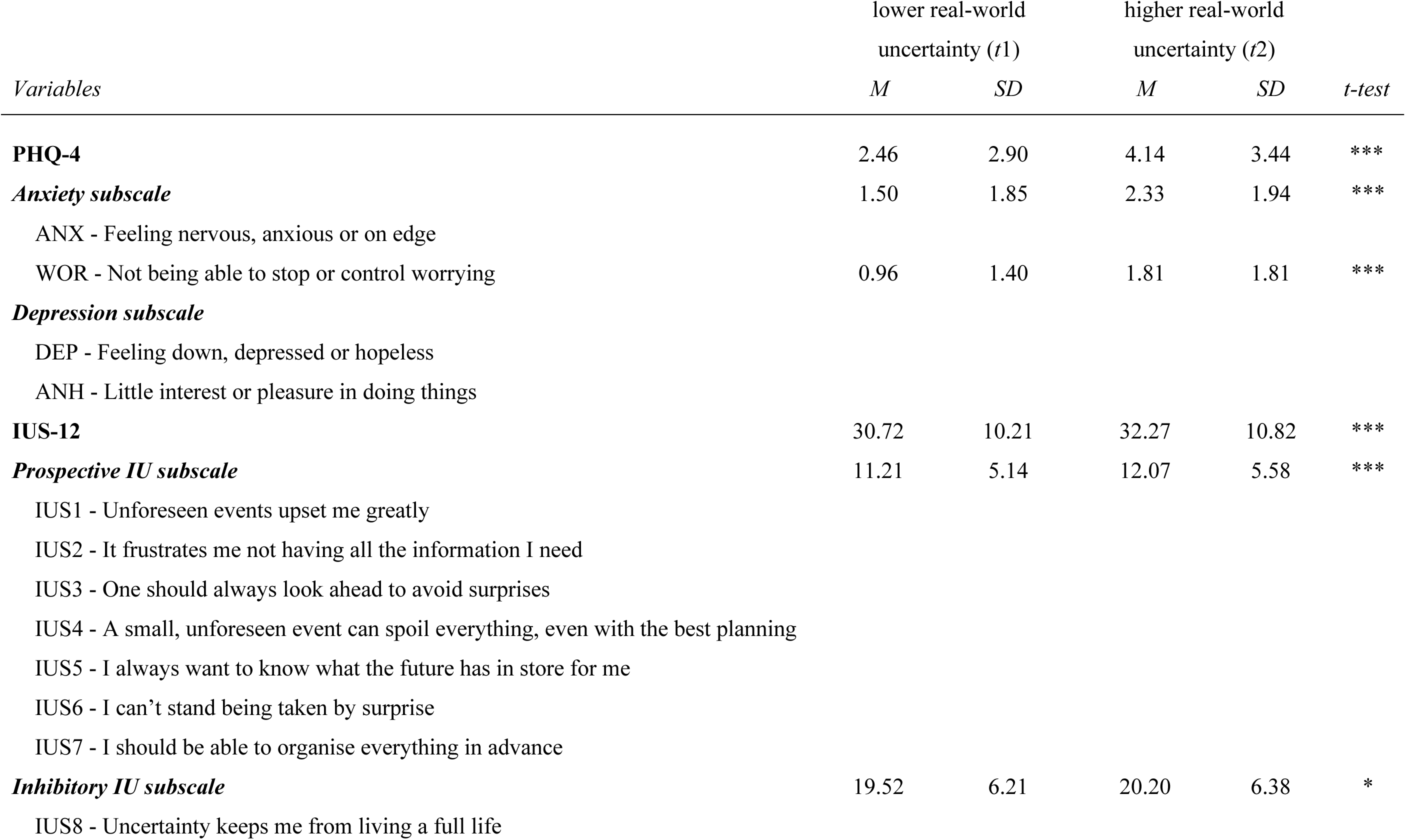
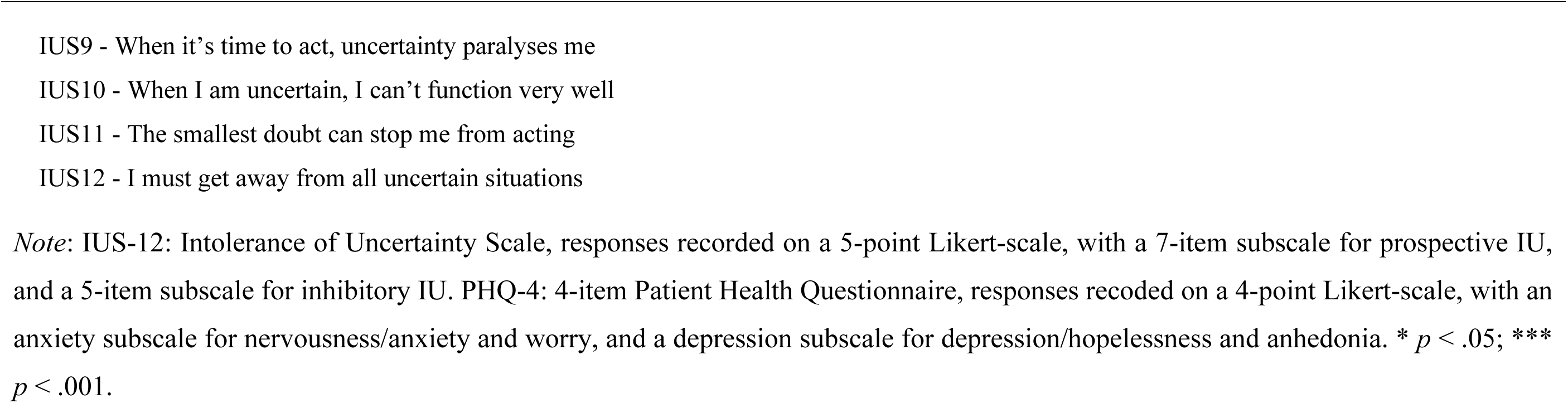
Symptom characteristics and list of measurement items.

### Data analysis

All data analyses were conducted in RStudio (R version 4.4.3).

#### Cross-sectional networks

To examine the structure of the relationship among variables, partial correlation network analysis was performed. Here, each node represents a variable (i.e., individual symptom/IU trait) and edges capture the relationships between nodes. In this context, a regularisation technique was used to penalise model complexity as to remove edges that are likely to be spurious, after controlling for all other variables (Epskamp et al., 2018; Foygel & Drton, 2010). Hence, edge weights represent the partial correlation coefficients between nodes, and variables independent from one another would have an edge weight of zero with no connection being drawn between these nodes. We first established regularised partial correlation networks based on standardised *z*-scores of the variables for both timepoints (i.e., lower and higher real-world uncertainty) via the *bootnet* package (Epskamp et al., 2018; Epskamp & Fried, 2021). Specifically, we estimated our networks through a least absolute shrinkage and selection operator (LASSO; Tibshirani (1996)), using the extended Bayesian information criterion (EBIC; Foygel and Drton (2010)) to estimate a sparse graphical model with a tuning hyperparameter (*γ*) of 0.5. This is a relatively stringent value of this parameter that minimises type-I errors and, at the expense of sensitivity, strengthens confidence in the emerging edges. This approach is recommended for polychoric correlation of polytomous data sets, like those obtained via questionnaires using Likert-scales in this study. We visualised the networks via the *qgraph* package (Epskamp et al., 2012).

##### Community detection

To establish communities (or clusters) of highly connected nodes, we used the Walktrap algorithm (Pons & Latapy, 2006) of the *igraph* package (Csardi & Nepusz, 2006; Csárdi et al., 2022). Here, communities are detected by performing random walks on a network and the partition with maximized modularity, as a measure of clustering strength, is subsequently selected. We then examined the modularity index *Q*, which provides a quantification of the intracommunity connectivity (Newman, 2004).

##### Centrality, accuracy and stability

We next assessed the centrality indices of node strength (i.e., sum of the absolute weights of every edge) and node bridge strength (i.e., sum of absolute weights of edges across clusters) using the *qgraph* and *networktools* packages (Epskamp et al., 2012; Jones, 2020). We did not report closeness and betweenness which have been shown to exhibit spurious associations among symptoms (Hallquist et al., 2021). For calibrated interpretations of network model results, it is important to examine accuracy measures by assessing fluctuations observed from sampling variability in edge weights. For each network model, we thus established the accuracy of the edge weights via non-parametric bootstraps with 2,000 samples each. The non-parametric bootstrapped resampling approach estimates confidence intervals (CIs) between the bootstrapped and sample means, with smaller and larger CIs suggestive of heightened and poorer precision of the edges, respectively. If a CI includes the value zero, this indicates that different node strengths (or edge weights) do not significantly differ from one another. We also assessed the stability of centrality estimates via case-dropping bootstraps with 2,000 samples each. The quantification of this stability was established using the correlation coefficients of stability (*CS*) which indicates whether centrality indices (i.e., strength and bridge strength) maintained stability when proportions of cases were progressively dropped (e.g., 10%, 20%, 30%). *CS* values stand for the maximum proportion of dropped samples, given that the correlation between original centrality indices could reach 0.7 at a 95% probability. A *CS* > .25 implies moderate stability and a *CS* > .5 implies strong stability (Epskamp et al., 2018).

#### Network comparison

Similarity of the two networks (lower and higher real-world uncertainty) was explored through correlations of the adjacency matrices and centrality indices (a value of 1 indicates perfect linear relationship, i.e., the same structure; a value of 0 indicates no relationship). Next, we used the NCT function of the *NetworkComparisonTest* package (van Borkulo et al., 2019) with 1,000 permutations and Bonferroni-Holm correction (van Borkulo et al., 2021). Network structure invariance, global strength (summed value of all network edges) and differences in edge weights were analysed.

#### Cross-lagged panel network (CLPN)

Finally, we conducted a two-step CLPN analysis of the data collected for lower (*t*1) and higher (*t*2) real-world environmental uncertainty (Wysocki et al., 2022). Central to CLPN modelling is the feature of directed paths across timepoints (in our example *t*1 and *t*2), reflecting connections among individual items (or themselves). These paths capture the variance shared between a variable at *t*1 and another (or the same) at *t*2, while all other variables are controlled for. In a first step, we included the estimation of regularised regression coefficients of each variable on itself (autoregressive paths) and on other variables (cross-lagged paths), from *t*1 to *t*2. The LASSO regression used draws on an *l*_1_ penalty via cross-validation, where small regression paths are shrunk to zero, leading to a sparse solution (Friedman et al., 2010). Nonetheless, LASSO regression biases the non-zero edges towards zero and the cross-validation technique has a high false positive rate to choose the penalty. Thus, in the second-step estimation, the model was re-estimated via a non-regularised regression approach. This was achieved by estimating the model selected in the first-step estimation within an SEM framework using the *lavaan* package (Rosseel, 2012). Here, the paths which were estimated to be zero in the regularised regression result (first-step estimation) were held zero. Finally, the model was further pruned by only using the significant parameters from non-regularised model (second-step estimation). We established centrality measures for cross-lagged in-strength and cross-lagged out-strength. Cross-lagged in-strength captures the predictability for each node and is calculated by summing the absolute weights of incoming edges. Cross-lagged out-strength captures the influence for each node, calculated by summing the absolute weights of outgoing edges.

## Results

### Descriptive statistics

Depression and anxiety symptoms and intolerance of uncertainty (i.e., PHQ-4 and IUS-12 scores) are shown in **Table 2**. Mean values of anxiety and depression symptoms were greater for *higher* (*t*2) compared to *lower* (*t*1) real-world uncertainty (*p* < .001). Furthermore, the proportion of participants who met the PHQ-4 subscales cut-off values drastically increased from lower to higher real-world uncertainty for anxiety (21.26% → 38.21%, *p* < .001) and depression (10.96% → 27.24%, *p* < .001). Pearsons correlations between the network variables and the two networks are graphically presented in **Figure 2a, b** and numerically in **Supplementary Table 1**-**2**.

**Figure 2.**
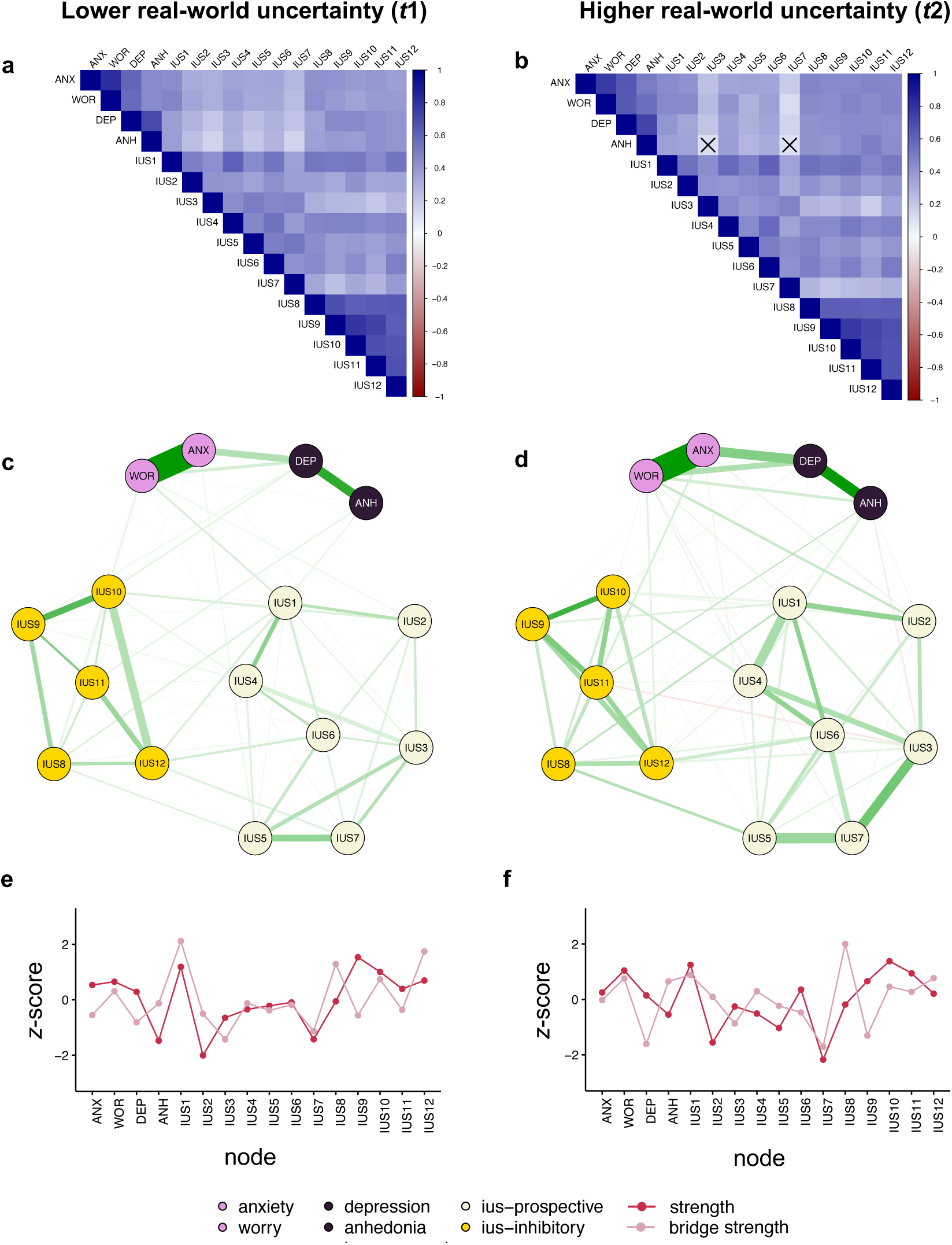
Correlations, cross-sectional networks, and centrality indices. **a** and **b** Pearson correlation of the measurements for lower real-world uncertainty (*t*1) and higher real-world uncertainty (*t*2). Darker blue reflects greater positive, darker red greater negative correlation. **c** and **d** Cross-sectional networks at *t*1 and *t*2. Green and red edges indicate positive and negative relationships, respectively. Edge thickness represents the strength with thicker edges indicating stronger relationships. Anxiety-related nodes are coloured in pink (ANX: anxiety/nervousness; WOR: worry), depression-related nodes in purple (DEP: depression/hopelessness; ANH: anhedonia), prospective IUS-subscale items in beige (IUS1-7), and inhibitory IUS-subscale items in gold (IUS8-12). IUS: Intolerance of Uncertainty Scale. **e** and **f** Centrality indices, strength (cherry red) and bridge strength (rose), reported in standardised *z*-scores with larger values reflecting greater centrality.

### Lower real-world uncertainty (*t*1) network

Three clusters (or communities) were identified, signalling coherent partitioning, with the first community including all four PHQ-items (ANX, WOR, DEP, ANH), the second community including the seven items of the prospective IUS subscale (IUS1-IUS7), and the third community including the five items of the inhibitory IUS subscale (IUS8-IUS12). The modularity of the clustering indicated moderate strength for the reported community structure (*Q* = .448). As shown in **Figure 2e**, nodes with the highest strength were IUS9, IUS1 and IUS10, while IUS1, IUS12 and IUS8 exhibited highest bridge strength. Descriptive values can be found in **Supplementary Table 3**. The coefficients of stability (*CS*) established via the case-dropping bootstrapping procedure revealed strong and moderate stability for node strength (*CS* = .518) and bridge strength (*CS* = .385), respectively. Accuracy of the edge weights, established via non-parametric bootstrapped, and correlation stability of centrality indices, are graphically represented in **Supplementary Figure 1a, c**.

### Higher real-world uncertainty (*t*2) network

For the higher real-world uncertainty (*t*2) network, the three clusters identified were identical to those reported for the lower real-world uncertainty (*t*1) network. The modularity of the clustering indicated a moderately strong community structure (*Q* = .415). As shown in **Figure 2f**, nodes with the highest strength were IUS10, IUS1 and WOR, while IUS8, IUS1 and IUS12 exhibited highest bridge strength. Descriptive values are reported in **Supplementary Table 3**. Node strength (*CS* = .449) and bridge strength (*CS* = .415) had moderate stability. Accuracy of the edge weights and correlation stability of centrality indices, are graphically represented in **Supplementary Figure 1b, d.**

### Network comparison

Comparing the two networks revealed linearity for edge weights and strong correlations for node strength and bridge strength (*rs* > .78, *ps* < .001, see **Table 3**). The Network Comparison Test showed no significant difference for network centrality indices and at the individual item level for centralities and strength.

**Table 3.**
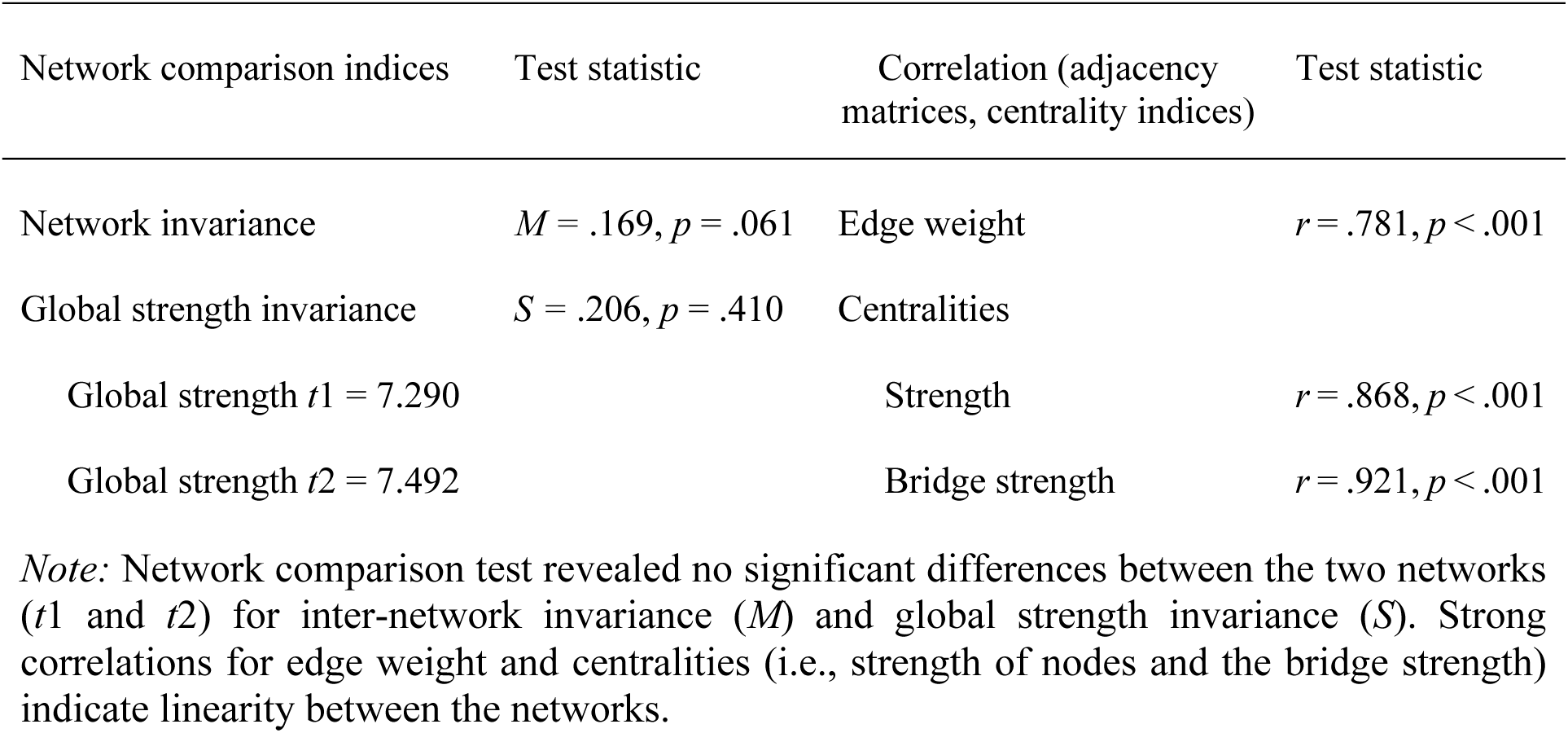
Network comparison (t1 to t2).

| Network comparison indices | Test statistic | Correlation (adjacency matrices, centrality indices) | Test statistic |
| --- | --- | --- | --- |
| Network invariance | $M = .169, p = .061$ | Edge weight | $r = .781, p < .001$ |
| Global strength invariance | $S = .206, p = .410$ | Centralities | |
| Global strength $t1 = 7.290$ | | Strength | $r = .868, p < .001$ |
| Global strength $t2 = 7.492$ | | Bridge strength | $r = .921, p < .001$ |

*Note:* Network comparison test revealed no significant differences between the two networks ( $t1$ and $t2$ ) for inter-network invariance ( $M$ ) and global strength invariance ( $S$ ). Strong correlations for edge weight and centralities (i.e., strength of nodes and the bridge strength) indicate linearity between the networks.

### Cross-lagged panel network (CLPN)

Results of the CLPN analysis are shown in **Figure 3a**, which includes both autoregressions and standardised between-nodes regressions (cross-lagged paths). The highest autoregression (see **Figure 3c**) values were observed for two prospective IU items (IUS5: *β* = .837; IUS6: *β* = .821) and two inhibitory IU items (IUS12: *β* = .813; IUS10: *β* = .794), while the lowest autoregression values were assigned to the four PHQ items (ANX: *β* = .289; ANH: *β* = .296; WOR: *β* = .343; DEP: *β* = .413). In other words, all autoregressive effects in IU outperformed those of anxiety and depression. To ease visual interpretation of the cross-lagged paths, we removed the autoregression and display the network graph in **Figure 3b**. The highest positive inter-node regressions were observed from IUS11 to anxiety (IUS11 → ANX: *β* = .221) and anhedonia to IUS2 (*β* = .164), highlighting that the item “The smallest doubt can stop me from acting*”* was the strongest predictor of anxiety and depression symptoms, following their autoregressive effects. The highest negative values were found for depression to IUS2 (*β* = -.179) and IUS3 to IUS8 (*β* = -.142). All values of the standardised regression matrix for the CPLN model are shown in **Supplementary Table 4.**

**Figure 3.**
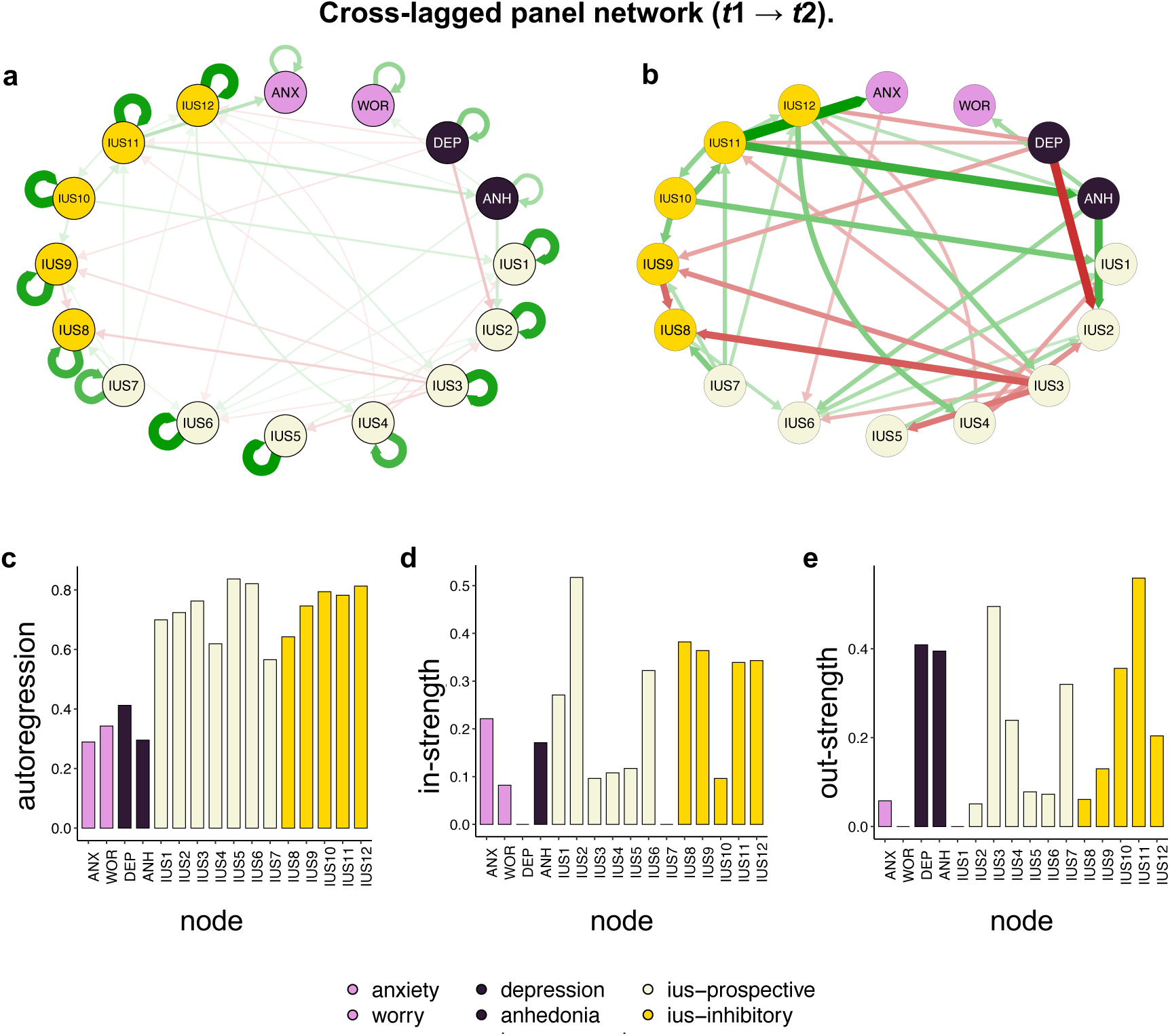
Cross-lagged panel network. **a** Full cross-lagged panel network (CLPN) with arrows representing unique longitudinal relationships. Green and red edges indicate positive and negative relationships, respectively. Edge thickness represents the strength with thicker edges indicating stronger relationships. Anxiety-related nodes are coloured in pink (ANX: anxiety/ nervousness; WOR: worry), depression-related nodes in purple (DEP: depression/ hopelessness; ANH: anhedonia), prospective IUS-subscale items in beige (IUS1-7), and inhibitory IUS-subscale items in gold (IUS8-12). IUS: Intolerance of Uncertainty Scale. **b** To ease visual interpretation of the between node relationships, autoregressive edges were removed from the full CLPN model. **c** Autoregression values of the individual measurement items. **d** Cross-lagged in-strength (i.e., sum of absolute weights of incoming edges) quantified the degree to which each variable (or node) at *t*2 was predicted by other variables (or nodes) at *t*1. **e** Cross-lagged out-strength (i.e., sum of absolute weights of outgoing edges) quantified the degree to which each variable (or node) at *t*1 predicted other variables (or nodes) at *t*2. *t*1: lower real-world uncertainty *t*2: higher real-world uncertainty.

Analysis of centralities showed that IUS items had the highest cross-lagged in-strength, meaning they were most highly predictable (IUS2 = .517; IUS8 = .382; IUS9 = .364, IUS12 = .343; see **Figure 3d**). Among the items with the highest cross-lagged out-strength, meaning most impacting other nodes were two IUS items and both depression items (IUS11 = .559; IUS3 = .495; DEP = 0.409; ANH = 0.395, see **Figure 3e**). Interestingly, while only negative inter-node regressions were reported for depression (DEP), only positive inter-node regressions were found for anhedonia (ANH). The complete set of in-strength and out-strength values are shown in **Supplementary Table 5.** Together, CLPN results highlighted that *behavioural inhibition due to doubt* (IUS11) positively predicted both anxiety and anhedonia. IUS items had the highest predictability and the highest autoregressive effects, indicating they were more stable than mental distress across varying levels of environmental uncertainty.

## Discussion

The current study established networks for intolerance of uncertainty (IU), anxiety and depression symptoms before and during a surge in real-world environmental uncertainty. We capitalised on data collected in the context of the COVID-19 pandemic–quantifiably differing in the World Uncertainty Index (**Figure 1a**)–as a naturalistic experiment of how global uncertainty interacts with IU to drive mental health symptoms. We also modelled directional longitudinal relationships between the symptoms of interest, allowing for temporal interpretations. Overall, our results show that worry became more central, and IU items served as strong autoregressive predictors during a period of heightened uncertainty, for which mental distress was elevated. Specifically, the symptom *inhibition of behaviour due to doubt* positively predicted anxiety and depression symptoms. This examination is of great relevance given the alarming prevalence rates of anxiety and depressive disorders amidst an increase of all-encompassing uncertainties worldwide. Furthermore, results of this study speak to the importance of how IU, as a proposed transdiagnostic factor, relates to internalising psychopathology symptoms in the face of uncertain environments.

In the lower real-world uncertainty context, the nodes with the highest strength and bridge strength belonged to the IU construct, while amidst higher real-world uncertainty, *worry* was among the highest strength nodes. Whereas times with comparably lower real-world uncertainty show lower symptom centrality, the centrality of IU indicates an increased vulnerability in those already concerned about potentially unexpected major events. This aligns with the notion of IU being a shared factor of emotional disorders (Gentes & Ruscio, 2011; McEvoy & Mahoney, 2011), meaning it exists, beyond a mere correlate, as a predisposition to negatively perceive and respond to uncertainty (Ladouceur et al., 1998; Ladouceur et al., 2000; Shihata et al., 2016). Among the anxiety and depression symptoms, *worry* emerged to play a more significant (central) role in times of heightened global uncertainty. As more central symptoms are thought to be crucial for disorder maintenance (Blanken et al., 2018; Bringmann et al., 2022; Rodebaugh et al., 2018), this finding further aligns with heightened occurrence of depression and generalised anxiety symptoms during the outbreak of the global pandemic (Hettich et al., 2022; McBride et al., 2021; van der Velden et al., 2021). Furthermore, establishing key symptoms and connecting symptoms in depressive-anxiety symptom network models has important clinical implications (Kaiser et al., 2021), as targeting these may enhance treatment effectiveness in patients and preventative care in high-risk individuals. Our findings highlight that during times of higher global uncertainty, the *cognitive* component of anxiety, that is anxious apprehension, operationalised as worry, becomes more relevant. Worry presents itself as a future-orientated, often uncontrollable, cognitive feature, with hyper-fixation on potential threat (Borkovec, 1994; Engels et al., 2007; Heller et al., 1997; Nitschke et al., 2001). We show that worry not only increases but also becomes more influential in the transdiagnostic symptom network during higher levels of real-world environmental uncertainty. Its heightened centrality suggests that worry develops increased influence on other nodes (e.g., depression or IU symptoms), potentially amplifying its effects within the overall network.

Directly comparing the cross-sectional networks, we found high correlations between network edges and centrality indices. This indicates that the network structures stayed intact when comparing the lower and higher uncertainty contexts, despite a worsening in symptom severity. Similarly, the high linearity between both uncertainty contexts and network invariance indices did not differ. Although a general trend of a higher strength for the higher compared to lower real-world uncertainty network was observable, this pattern did not reach statistical significance, which may be influenced by a lack of sufficient power. Previous work with a highly powered sample size has highlighted how IU is an important risk factor for internalising psychopathology during heightened global uncertainty (Andrews et al., 2023). Future research may examine longitudinal differences across distinct periods of higher and lower uncertainty in larger sample sizes.

Using a community detection algorithm, we identified three communities for both cross-sectional networks, which corresponded with the entire set of anxiety and depression symptoms, as well as the two domains of the Intolerance of Uncertainty Scale (IUS; inhibitory-IU and prospective-IU subscale). In other words, the algorithm did not establish separate communities for anxiety and depression symptoms. This not only validates the factor structure of the IUS but also highlights the close interconnection and comorbidity of anxiety and depression symptomatology (Carleton et al., 2007; Kroenke et al., 2009; Löwe et al., 2010). In the context of recent developments towards empirically-based research frameworks and classification of mental disorders (e.g., Research Domain Criteria (Insel et al., 2010), and Hierarchical Taxonomy of Psychopathology (Kotov et al., 2017)) as well as computational modelling approaches (Adams et al., 2016; Hauser et al., 2022; Lawson et al., 2021; Stephan & Mathys, 2014; Zhang et al., 2025), our findings reinforce the value of transdiagnostic approaches that focus on symptom-level interactions rather than traditional diagnostic boundaries. The overlapping community structure observed in our networks reflects the empirical reality of shared mechanisms and symptom co-occurrence across disorders, supporting efforts to refine classification systems and tailor interventions to underlying processes rather than diagnostic labels. However, future work should test the robustness of this community structure using more granular and comprehensive measures of anxiety and depression symptoms.

Results from the cross-lagged panel network show that autoregressions of the IU items were higher than anxiety and depression symptoms. Autoregressive paths highlight the stability of individual items between the timepoints of measure, reflecting idiosyncratic impacts across *t*1 and *t*2. In other words, IU variables outperformed internalising psychopathology variables in their value in predicting anxiety and depression symptoms. IU has been conceptualised as a trait-like, trans-situational variable (Buhr & Dugas, 2002). Thus, this personality-type feature may hold important, yet overlooked, predictive value beyond the traditional psychopathological symptoms (e.g., anxiety and depression) in informing the onset and maintenance of mental health difficulties. Our results show that this holds especially true in the face of increased real-world uncertainty. Indeed, in the context of the all-encompassing uncertain environments many individuals now face, our findings underscore the importance of targeting IU in both preventative and therapeutic interventions, and of using it as a marker of vulnerability before major stressors occur (Boswell et al., 2013; Carleton, 2016; McEvoy et al., 2019). In line with recent theoretical work highlighting IU as a candidate mechanism for the development, persistence, and treatment of mood and anxiety disorders (Sandhu et al., 2023), our findings further support positioning IU at the centre of transdiagnostic models of internalising psychopathology.

Furthermore, notable relationships between individual symptoms emerged in the longitudinal modelling reported in this study. A key aspect of interpreting network paths in the CLPN is that they reflect conditional predictive relationships–that is, the influence of each variable at *t*1 on *t*2, while controlling for all other variables at *t*1. One particularly salient finding was that *inhibition of behaviour due to doubt* (IUS11) at times of lower uncertainty (*t*1), positively predicted anxiety (ANX) and anhedonia (ANH) during higher uncertainty (*t*2). This aligns with previous research implicating behavioural inhibition as a risk factor for anxiety (Svihra & Katzman, 2004) and with neurocognitive models emphasising behavioural and cognitive avoidance as central mechanisms of anxiety under uncertainty (Grupe & Nitschke, 2013). Our findings suggest that behavioural inhibition rooted in doubt may represent a treatment target in both cognitive behavioural therapy and preventative mental health strategies.

Within cognitive therapy approaches for anxiety disorders, there is emphasis on attention bias modification (ABM, Mogg and Bradley (2016); Mogg et al. (2017); Pergamin-Hight et al. (2015)). These methods support goal-engagement factors in the context of attention but could be extended to disengaging individuals from maladaptive doubts. Similarly, by promoting acceptance, mindfulness, values, and committed action, therapy approaches such as Acceptance and Commitment Therapy (ACT, Hayes et al. (2012); Roemer and Orsillo (2008)) help individuals build psychological flexibility. This flexibility is crucial for managing self-doubt and taking steps towards a more fulfilling life, making ACT a particularly promising treatment approach for uncertain situations.

Interestingly, we also found that anhedonia (ANH) positively predicted *frustration of not having relevant information* (IUS2), whereas depression (DEP) predicted the same item negatively. Individuals who reported high levels of anhedonia were more frustrated in situations involving a lack of relevant information; at the same time, individuals who reported high levels of depression were less frustrated in these contexts. This distinction aligns with theoretical models that differentiate motivational difficulties (such as reduced approach behaviour and reward sensitivity in anhedonia) from the broader cognitive and affective profile of depression, which may include hopelessness, resignation, and reduced cognitive engagement with the external world (Beck, 1976; Huys et al., 2015; Treadway & Zald, 2011). Individuals high in anhedonia may still exhibit a desire for goal-directed information, despite blunted hedonic responsiveness—and may feel more acutely frustrated when such information is unavailable. Conversely, individuals with more pervasive depressive symptoms may be less responsive to uncertainty due to learned helplessness. This dissociation suggests that frustration in uncertain contexts may reflect a residual or thwarted form of goal pursuit, preserved in some internalising profiles but diminished in others.

## Limitations

The findings of this network analysis study should be interpreted in the context of its limitations. First, we reported data collected from a community (opportunity) sample, an approach that enhances generalisability, but may be criticised for its lack to account for interaction of transdiagnostic symptoms in a clinical population. However, around a third of participants reached cut-off values for both depression and anxiety symptoms in the context of higher real-world uncertainty, highlighting the presence of clinically significant levels of psychopathology in our sample. Relatedly, all data were based on self-report measures rather than formal clinical assessments. In light of ongoing debates about optimal approaches to psychopathology assessment (American Psychiatric Association DSM-5 Task Force, 2013; Insel et al., 2010; Kotov et al., 2017), future studies should integrate categorical, dimensional, and transdiagnostic methods—including diagnostic interviews—to provide a more comprehensive picture. Specifically, cross-sectional and temporal network analyses in clinical cohorts with anxiety and/or depressive disorders would help assess the robustness and clinical significance of the findings.

Second, participants reported on both timepoints—representing periods of differing levels of real-world uncertainty—concurrently. This retrospective reporting introduces the possibility of recall bias (Brewin et al., 1993; Solhan et al., 2009), particularly given the emotional salience and duration of the COVID-19 pandemic. Although baseline data were unavailable due to the unanticipated onset of the pandemic, retrospective self-report remains common in psychological and clinical research. Instruments such as the STAI-T, IUS, and the PBI routinely ask individuals to reflect on their *typical* behaviour or emotional experiences, or to recall earlier life periods. While such reports may be influenced by current mood or context, they nonetheless offer valid and meaningful insights into perceived patterns of functioning over time (Achenbach et al., 2005).

Third, there are ongoing methodological challenges and unresolved assumptions in psychological network modelling (Pearl, 2000, 2012). Although longitudinal network modelling can reveal the direction of potentially causal relationships, it is debated what time lags are appropriate or optimal to capture relationships between symptoms (Jacobson et al., 2019; Wysocki et al., 2022). Indeed, relationships between symptoms may occur at shorter or longer time periods, as parameter estimates vary as a function of the time interval between measurements (Gollob & Reichardt, 1987). To address this, future research could employ high-frequency sampling techniques—such as daily diary methods or ecological momentary assessment (EMA)—to track real-time fluctuations in IU and emotional responses to environmental stressors. These approaches would offer a more granular and temporally sensitive perspective on how IU dynamically interacts with real-world uncertainty, potentially yielding more causally informative insights than static, retrospective survey measures.

## Conclusion

Employing network analyses on transdiagnostic symptomatology, we investigated how real-world environmental uncertainty interacts with intolerance of uncertainty (IU) to drive mental health symptoms. Cross-sectional networks highlighted that worry became more central in times of higher real-world uncertainty, while cross-lagged panel network modelling revealed that inhibition of behaviour due to doubt, reported during times of relative global stability, positively predicted both anxiety and anhedonia. Notably, autoregressive effects suggest that during periods of greater certainty, internalising symptoms are poorer predictors of future distress. In contrast, transdiagnostic traits like IU are more stable and, crucially, serve as predictors of symptoms and mental distress, even when distress itself is not necessarily present. This suggests that trait-level cognitive vulnerabilities, such as IU, can anticipate the onset or worsening of symptoms—even when overt distress is not yet observable. In an era defined by global and personal uncertainty, these findings offer new insights into who is vulnerable, when, and why. By identifying IU-related processes as both persistent and predictive, our results underscore the value of targeting these mechanisms in early intervention and treatment.

## References

Achenbach, T. M., Krukowski, R. A., Dumenci, L., & Ivanova, M. Y. (2005). Assessment of adult psychopathology: meta-analyses and implications of cross-informant correlations. Psychological Bulletin, 131(3), 361.

Adams, R. A., Huys, Q. J., & Roiser, J. P. (2016). Computational psychiatry: towards a mathematically informed understanding of mental illness. Journal of Neurology, Neurosurgery and Psychiatry, 87(1), 53–63.

Ahir, H., Bloom, N., & Furceri, D. (2022). The World Uncertainty Index. National Bureau of Economic Research Working Paper Series, No. 29763. 10.3386/w29763

Altig, D., Baker, S., Barrero, J. M., Bloom, N., Bunn, P., Chen, S., Davis, S. J., Leather, J., Meyer, B., & Mihaylov, E. (2020). Economic uncertainty before and during the COVID-19 pandemic. Journal of public economics, 191, 104274.

American Psychiatric Association DSM-5 Task Force. (2013). Diagnostic and statistical manual of mental disorders: DSM-5™, 5th ed. American Psychiatric Publishing, Inc. 10.1176/appi.books.9780890425596

Andrews, J. L., Li, M., Minihan, S., Songco, A., Fox, E., Ladouceur, C. D., Mewton, L., Moulds, M., Pfeifer, J. H., Van Harmelen, A. L., & Schweizer, S. (2023). The effect of intolerance of uncertainty on anxiety and depression, and their symptom networks, during the COVID-19 pandemic. BMC Psychiatry, 23(1), 261. 10.1186/s12888-023-04734-8

Beck, A. T. (1976). Cognitive therapy and the emotional disorders. International Universities Press, Oxford.

Blanken, T. F., Deserno, M. K., Dalege, J., Borsboom, D., Blanken, P., Kerkhof, G. A., & Cramer, A. O. (2018). The role of stabilizing and communicating symptoms given overlapping communities in psychopathology networks. Scientific Reports, 8(1), 5854.

Borkovec, T. D. (1994). The nature, functions, and origins of worry. In Worrying: Perspectives on theory, assessment and treatment. (pp. 5–33). John Wiley & Sons.

Borsboom, D. (2017). A network theory of mental disorders. World Psychiatry, 16, 5–13. 10.1002/wps.20375

Boswell, J. F., Thompson-Hollands, J., Farchione, T. J., & Barlow, D. H. (2013). Intolerance of uncertainty: a common factor in the treatment of emotional disorders. Journal of Clinical Psychology, 69(6), 630–645. 10.1002/jclp.21965

Bower, G. H. (1987). Commentary on mood and memory. Behaviour Research and Therapy, 25(6), 443–455.

Brewin, C. R., Andrews, B., & Gotlib, I. H. (1993). Psychopathology and early experience: a reappraisal of retrospective reports. Psychological Bulletin, 113(1), 82.

Bringmann, L. F., Albers, C., Bockting, C., Borsboom, D., Ceulemans, E., Cramer, A., Epskamp, S., Eronen, M. I., Hamaker, E., Kuppens, P., Lutz, W., McNally, R. J., Molenaar, P., Tio, P., Voelkle, M. C., & Wichers, M. (2022). Psychopathological networks: Theory, methods and practice. Behaviour Research and Therapy, 149, 104011. 10.1016/j.brat.2021.104011

Buhr, K., & Dugas, M. J. (2002). The Intolerance of Uncertainty Scale: psychometric properties of the English version. Behaviour Research and Therarpy, 40(8), 931–945. 10.1016/s0005-7967(01)00092-4

Carleton, R. N. (2016). Into the unknown: A review and synthesis of contemporary models involving uncertainty. Journal of Anxiety Disorders, 39, 30–43. 10.1016/j.janxdis.2016.02.007

Carleton, R. N., Norton, M. P. J., & Asmundson, G. J. (2007). Fearing the unknown: A short version of the Intolerance of Uncertainty Scale. Journal of Anxiety Disorders, 21(1), 105–117.

Coughlin, S. S. (1990). Recall bias in epidemiologic studies. Journal of Clinical Epidemiology, 43(1), 87–91. 10.1016/0895-4356(90)90060-3

Cramer, A., Waldorp, L., Maas, H., & Borsboom, D. (2010). Comorbidity: A network perspective. The Behavioral and brain sciences, 33, 137–150; discussion 150. 10.1017/S0140525X09991567

Csardi, G., & Nepusz, T. (2006). The igraph software package for complex network research. Complex Systems, 1695, 1–9.

Csárdi, G., Nepusz, T., Traag, V., Horvát, S., Zanini, F., Noom, D., & Müller, K. (2022). igraph: Network analysis and visualization in R (2.1.1). R-CRAN. https://CRAN.R-project.org/package=igraph

Dugas, M. J., Buhr, K., & Ladouceur, R. (2004). The Role of Intolerance of Uncertainty in Etiology and Maintenance. In R. G. Heimberg (Ed.), Generalized Anxiety Disorder: Advances in Research And Practice (pp. 143–163). The Guilford Press.

Dugas, M. J., Gagnon, F., Ladouceur, R., & Freeston, M. H. (1998). Generalized anxiety disorder: a preliminary test of a conceptual model. Behaviour Research and Therapy, 36(2), 215–226. 10.1016/s0005-7967(97)00070-3

Dugas, M. J., Sexton, K. A., Hebert, E. A., Bouchard, S., Gouin, J.-P., & Shafran, R. (2022). Behavioral Experiments for Intolerance of Uncertainty: A Randomized Clinical Trial for Adults With Generalized Anxiety Disorder. Behavior Therapy, 53(6), 1147–1160. 10.1016/j.beth.2022.05.003

Engels, A. S., Heller, W., Mohanty, A., Herrington, J. D., Banich, M. T., Webb, A. G., & Miller, G. A. (2007). Specificity of regional brain activity in anxiety types during emotion processing. Psychophysiology, 44(3), 352–363.

Epskamp, S., Borsboom, D., & Fried, E. I. (2018). Estimating psychological networks and their accuracy: A tutorial paper. Behavior Research Methods, 50(1), 195–212. 10.3758/s13428-017-0862-1

Epskamp, S., Cramer, A. O. J., Waldorp, L. J., Schmittmann, V. D., & Borsboom, D. (2012). qgraph: Network Visualizations of Relationships in Psychometric Data. Journal of Statistical Software, 48(4), 1–18. 10.18637/jss.v048.i04

Epskamp, S., & Fried, E. (2021). bootnet: Bootstrap methods for various network estimation routines (1.6). R-CRAN. https://cran.r-project.org/package=bootnet

Faul, L., & LaBar, K. S. (2023). Mood-congruent memory revisited. Psychological Review, 130(6), 1421–1456. 10.1037/rev0000394

Foygel, R., & Drton, M. (2010). Extended Bayesian information criteria for Gaussian graphical models. Advances in Neural Information Processing Systems, 23.

Friedman, J., Hastie, T., & Tibshirani, R. (2010). Regularization paths for generalized linear models via coordinate descent. Journal of Statistical Software, 33(1), 1.

Gentes, E. L., & Ruscio, A. M. (2011). A meta-analysis of the relation of intolerance of uncertainty to symptoms of generalized anxiety disorder, major depressive disorder, and obsessive–compulsive disorder. Clinical Psychology Review, 31(6), 923–933.

Global Burden of Disease Collaborative Network. Global Burden of Disease Study 2021 (GBD 2021). In. Seattle, United States: Institute for Health Metrics and Evaluation (IHME), 2024.

Gollob, H. F., & Reichardt, C. S. (1987). Taking account of time lags in causal models. Child Development, 58(1), 80–92. 10.2307/1130293

Groen, R. N., Wichers, M., Wigman, J. T. W., & Hartman, C. A. (2019). Specificity of psychopathology across levels of severity: a transdiagnostic network analysis. Scientific Reports, 9(1), 18298. 10.1038/s41598-019-54801-y

Grupe, D. W., & Nitschke, J. B. (2013). Uncertainty and anticipation in anxiety: an integrated neurobiological and psychological perspective. Nature Reviews Neuroscience, 14(7), 488–501.

Hallquist, M. N., Wright, A. G. C., & Molenaar, P. C. M. (2021). Problems with Centrality Measures in Psychopathology Symptom Networks: Why Network Psychometrics Cannot Escape Psychometric Theory. Multivariate Behav Res, 56(2), 199–223. 10.1080/00273171.2019.1640103

Harris, P. A., Taylor, R., Minor, B. L., Elliott, V., Fernandez, M., O’Neal, L., McLeod, L., Delacqua, G., Delacqua, F., Kirby, J., & Duda, S. N. (2019). The REDCap consortium: Building an international community of software platform partners. Journal of Biomedical Informatics, 95, 103208. 10.1016/j.jbi.2019.103208

Harris, P. A., Taylor, R., Thielke, R., Payne, J., Gonzalez, N., & Conde, J. G. (2009). Research electronic data capture (REDCap)--a metadata-driven methodology and workflow process for providing translational research informatics support. Journal of Biomedical Informatics, 42(2), 377–381. 10.1016/j.jbi.2008.08.010

Hauser, T. U., Skvortsova, V., De Choudhury, M., & Koutsouleris, N. (2022). The promise of a model-based psychiatry: building computational models of mental ill health. The Lancet Digital Health, 4(11), e816–e828.

Hayes, S. C., Strosahl, K. D., & Wilson, K. G. (2012). Acceptance and commitment therapy: The process and practice of mindful change, 2nd ed. The Guilford Press.

Hedley, F. E., Larsen, E., Mohanty, A., Liu, J. Z., & Jin, J. (2024). Understanding anxiety through uncertainty quantification. British Journal of Psychology, 00, 1–14. 10.1111/bjop.12693

Heller, W., Nitschke, J. B., Etienne, M. A., & Miller, G. A. (1997). Patterns of regional brain activity differentiate types of anxiety. Journal of Abnormal Psychology, 106(3), 376.

Hertz-Palmor, N., Moore, T. M., Gothelf, D., DiDomenico, G. E., Dekel, I., Greenberg, D. M., Brown, L. A., Matalon, N., Visoki, E., White, L. K., Himes, M. M., Schwartz-Lifshitz, M., Gross, R., Gur, R. C., Gur, R. E., Pessach, I. M., & Barzilay, R. (2021). Association among income loss, financial strain and depressive symptoms during COVID-19: Evidence from two longitudinal studies. Journal of Affective Disorders, 291, 1–8. 10.1016/j.jad.2021.04.054

Hertz-Palmor, N., Ruppin, S., Matalon, N., Mosheva, M., Dorman-Ilan, S., Serur, Y., Avinir, A., Mekori-Domachevsky, E., Hasson-Ohayon, I., Gross, R., Gothelf, D., & Pessach, I. M. (2023). A 16-month longitudinal investigation of risk and protective factors for mental health outcomes throughout three national lockdowns and a mass vaccination campaign: Evidence from a weighted Israeli sample during COVID-19. Psychiatry Research, 323, 115119. 10.1016/j.psychres.2023.115119

Hettich, N., Entringer, T. M., Kroeger, H., Schmidt, P., Tibubos, A. N., Braehler, E., & Beutel, M. E. (2022). Impact of the COVID-19 pandemic on depression, anxiety, loneliness, and satisfaction in the German general population: a longitudinal analysis. Social Psychiatry and Psychiatric Epidemiology, 57(12), 2481–2490. 10.1007/s00127-022-02311-0

Huys, Q. J. M., Daw, N. D., & Dayan, P. (2015). Depression: A Decision-Theoretic Analysis. Annual Review of Neuroscience, 38(Volume 38, 2015), 1-23. 10.1146/annurev-neuro-071714-033928

Insel, T., Cuthbert, B., Garvey, M., Heinssen, R., Pine, D. S., Quinn, K., Sanislow, C., & Wang, P. (2010). Research Domain Criteria (RDoC): Toward a New Classification Framework for Research on Mental Disorders. The American Journal of Psychiatry 167(7), 748–751. 10.1176/appi.ajp.2010.09091379

Jach, H. K., & Smillie, L. D. (2019). To fear or fly to the unknown: Tolerance for ambiguity and Big Five personality traits. Journal of Research in Personality, 79, 67–78. 10.1016/j.jrp.2019.02.003

Jacobson, N. C., Chow, S.-M., & Newman, M. G. (2019). The Differential Time-Varying Effect Model (DTVEM): A tool for diagnosing and modeling time lags in intensive longitudinal data. Behavior Research Methods, 51(1), 295–315. 10.3758/s13428-018-1101-0

Jones, P. (2020). Networktools: Tools for identifying important nodes in networks (1.5.2). R-CRAN. https://cran.r-project.org/package=networktools

Kaiser, T., Herzog, P., Voderholzer, U., & Brakemeier, E. L. (2021). Unraveling the comorbidity of depression and anxiety in a large inpatient sample: Network analysis to examine bridge symptoms. Depression and Anxiety, 38(3), 307–317. 10.1002/da.23136

Kessler, R. C., Chiu, W. T., Demler, O., & Walters, E. E. (2005). Prevalence, severity, and comorbidity of 12-month DSM-IV disorders in the National Comorbidity Survey Replication. Archives of General Psychiatry, 62(6), 617–627.

Kienzler, H., Giacaman, R., & Massazza, A. (2022). The association between uncertainty and mental health: a scoping review of the quantitative literature. Journal of Mental Health, 32. 10.1080/09638237.2021.2022620

Kotov, R., Krueger, R. F., Watson, D., Achenbach, T. M., Althoff, R. R., Bagby, R. M., Brown, T. A., Carpenter, W. T., Caspi, A., Clark, L. A., Eaton, N. R., Forbes, M. K., Forbush, K. T., Goldberg, D., Hasin, D., Hyman, S. E., Ivanova, M. Y., Lynam, D. R., Markon, K., . . . Zimmerman, M. (2017). The Hierarchical Taxonomy of Psychopathology (HiTOP): A dimensional alternative to traditional nosologies. Journal of Abnormal Psychology, 126(4), 454–477. 10.1037/abn0000258

Kroenke, K., Spitzer, R. L., Williams, J. B. W., & Löwe, B. (2009). An Ultra-Brief Screening Scale for Anxiety and Depression: The PHQ–4. Psychosomatics, 50(6), 613–621. 10.1016/S0033-3182(09)70864-3

Ladouceur, R., Blais, F., Freeston, M. H., & Dugas, M. J. (1998). Problem solving and problem orientation in generalized anxiety disorder. Journal of Anxiety Disorders, 12(2), 139–152.

Ladouceur, R., Gosselin, P., & Dugas, M. J. (2000). Experimental manipulation of intolerance of uncertainty: A study of a theoretical model of worry. Behaviour Research and Therapy, 38(9), 933–941.

Lawson, R. P., Bisby, J., Nord, C. L., Burgess, N., & Rees, G. (2021). The Computational, Pharmacological, and Physiological Determinants of Sensory Learning under Uncertainty. Current Biology, 31(1), 163–172.e164. 10.1016/j.cub.2020.10.043

Löwe, B., Wahl, I., Rose, M., Spitzer, C., Glaesmer, H., Wingenfeld, K., Schneider, A., & Brähler, E. (2010). A 4-item measure of depression and anxiety: Validation and standardization of the Patient Health Questionnaire-4 (PHQ-4) in the general population. Journal of Affective Disorders, 122(1), 86–95. 10.1016/j.jad.2009.06.019

McBride, O., Murphy, J., Gibson-Miller, J., Hartman, T., Hyland, P., Levita, L., Mason, L., Martinez, A., McKay, R., Stocks, T., Bennett, K., Vallières, F., Karatzias, T., Valiente, C., Vazquez, C., & Bentall, R. (2020). Monitoring the psychological, social, and economic impact of the COVID-19 pandemic in the population: Context, design and conduct of the longitudinal COVID-19 psychological research consortium (C19PRC) study. International Journal of Methods in Psychiatric Research, 30. 10.1002/mpr.1861

McBride, O., Murphy, J., Shevlin, M., Gibson-Miller, J., Hartman, T. K., Hyland, P., Levita, L., Mason, L., Martinez, A. P., McKay, R., Stocks, T. V., Bennett, K. M., Vallières, F., Karatzias, T., Valiente, C., Vazquez, C., & Bentall, R. P. (2021). Monitoring the psychological, social, and economic impact of the COVID-19 pandemic in the population: Context, design and conduct of the longitudinal COVID-19 psychological research consortium (C19PRC) study. International Journal of Methods in Psychiatric Research, 30(1), e1861. 10.1002/mpr.1861

McEvoy, P. M., Hyett, M. P., Shihata, S., Price, J. E., & Strachan, L. (2019). The impact of methodological and measurement factors on transdiagnostic associations with intolerance of uncertainty: A meta-analysis. Clinical Psychology Review, 73, 101778.

McEvoy, P. M., & Mahoney, A. E. (2011). Achieving certainty about the structure of intolerance of uncertainty in a treatment-seeking sample with anxiety and depression. Journal of Anxiety Disorders, 25(1), 112–122.

Moffitt, T. E., Caspi, A., Taylor, A., Kokaua, J., Milne, B. J., Polanczyk, G., & Poulton, R. (2010). How common are common mental disorders? Evidence that lifetime prevalence rates are doubled by prospective versus retrospective ascertainment. Psychological Medicine, 40(6), 899–909. 10.1017/S0033291709991036

Mogg, K., & Bradley, B. P. (2016). Anxiety and attention to threat: Cognitive mechanisms and treatment with attention bias modification. Behaviour Research and Therapy, 87, 76–108. 10.1016/j.brat.2016.08.001

Mogg, K., Waters, A. M., & Bradley, B. P. (2017). Attention Bias Modification (ABM): Review of Effects of Multisession ABM Training on Anxiety and Threat-Related Attention in High-Anxious Individuals. Clin Psychol Sci, 5(4), 698–717. 10.1177/2167702617696359

Newman, M. E. (2004). Analysis of weighted networks. Physical Review E—Statistical, Nonlinear, and Soft Matter Physics, 70(5). 10.1103/PhysRevE.70.056131

Newman, M. E. J. (2006). Modularity and community structure in networks. Proceedings of the National Academy of Sciences, 103(23), 8577–8582. doi:10.1073/pnas.0601602103

Nitschke, J. B., Heller, W., Imig, J. C., McDonald, R. P., & Miller, G. A. (2001). Distinguishing Dimensions of Anxiety and Depression. Cognitive Therapy and Research, 25(1), 1–22. 10.1023/A:1026485530405

Pearl, J. (2000). Causality: Models, reasoning, and inference. Cambridge University Press.

Pearl, J. (2012). The causal mediation formula--a guide to the assessment of pathways and mechanisms. Prev Sci, 13(4), 426–436. 10.1007/s11121-011-0270-1

Pergamin-Hight, L., Naim, R., Bakermans-Kranenburg, M. J., van, I. M. H., & Bar-Haim, Y. (2015). Content specificity of attention bias to threat in anxiety disorders: a meta-analysis. Clinical Psychology Review, 35, 10–18. 10.1016/j.cpr.2014.10.005

Ramos-Vera, C., García O’Diana, A., Basauri-Delgado, M., Calizaya-Milla, Y. E., & Saintila, J. (2024). Network analysis of anxiety and depressive symptoms during the COVID-19 pandemic in older adults in the United Kingdom. Scientific Reports, 14(1), 7741. 10.1038/s41598-024-58256-8

Ren, L., Yang, Z., Wang, Y., Cui, L.-B., Jin, Y., Ma, Z., Zhang, Q., Wu, Z., Wang, H.-N., & Yang, Q. (2020). The relations among worry, meta-worry, intolerance of uncertainty and attentional bias for threat in men at high risk for generalized anxiety disorder: a network analysis. BMC Psychiatry, 20(1), 452. 10.1186/s12888-020-02849-w

Robichaud, M. (2013). Cognitive Behavior Therapy Targeting Intolerance of Uncertainty: Application to a Clinical Case of Generalized Anxiety Disorder. Cognitive and Behavioral Practice, 20(3), 251–263. 10.1016/j.cbpra.2012.09.001

Rodebaugh, T. L., Tonge, N. A., Piccirillo, M. L., Fried, E., Horenstein, A., Morrison, A. S., Goldin, P., Gross, J. J., Lim, M. H., & Fernandez, K. C. (2018). Does centrality in a cross-sectional network suggest intervention targets for social anxiety disorder? Journal of Consulting and Clinical Psychology, 86(10), 831.

Roemer, L., & Orsillo, S. M. (2008). Mindfulness-and acceptance-based behavioral therapies in practice. Guilford Press.

Rosseel, Y. (2012). lavaan: An R package for structural equation modeling. Journal of Statistical Software, 48, 1–36.

Rosser, B. A. (2019). Intolerance of Uncertainty as a Transdiagnostic Mechanism of Psychological Difficulties: A Systematic Review of Evidence Pertaining to Causality and Temporal Precedence. Cognitive Therapy and Research, 43(2), 438–463. 10.1007/s10608-018-9964-z

Sandhu, T. R., Xiao, B., & Lawson, R. P. (2023). Transdiagnostic computations of uncertainty: towards a new lens on intolerance of uncertainty. Neuroscience and Biobehavioral Reviews, 148, 105123. 10.1016/j.neubiorev.2023.105123

Santomauro, D. F., Herrera, A. M. M., Shadid, J., Zheng, P., Ashbaugh, C., Pigott, D. M., Abbafati, C., Adolph, C., Amlag, J. O., & Aravkin, A. Y. (2021). Global prevalence and burden of depressive and anxiety disorders in 204 countries and territories in 2020 due to the COVID-19 pandemic. The Lancet, 398(10312), 1700–1712.

Schweizer, S., Lawson, R., & Blakemore, S.-J. (2023). Uncertainty as a driver of the youth mental health crisis. Current Opinion in Psychology, 53, 101657. 10.1016/j.copsyc.2023.101657

Shihata, S., McEvoy, P. M., Mullan, B. A., & Carleton, R. N. (2016). Intolerance of uncertainty in emotional disorders: What uncertainties remain? Journal of Anxiety Disorders, 41, 115–124.

Solhan, M. B., Trull, T. J., Jahng, S., & Wood, P. K. (2009). Clinical assessment of affective instability: Comparing EMA indices, questionnaire reports, and retrospective recall. Psychological Assessment, 21(3), 425–436. 10.1037/a0016869

Stephan, K. E., & Mathys, C. (2014). Computational approaches to psychiatry. Current Opinion in Neurobiology, 25, 85–92.

Svihra, M., & Katzman, M. A. (2004). Behavioural inhibition: A predictor of anxiety. Paediatrics & Child Health, 9(8), 547–550. 10.1093/pch/9.8.547

Talari, K., & Goyal, M. (2020). Retrospective studies–utility and caveats. Journal of the Royal College of Physicians of Edinburgh, 50(4), 398–402.

Tibshirani, R. (1996). Regression Shrinkage and Selection via the Lasso. Journal of the Royal Statistical Society. Series B (Methodological), 58(1), 267–288. http://www.jstor.org/stable/2346178

Treadway, M. T., & Zald, D. H. (2011). Reconsidering anhedonia in depression: lessons from translational neuroscience. Neuroscience and Biobehavioral Reviews, 35(3), 537–555.

van Borkulo, C., Epskamp, S., & Jones, P. (2019). NetworkComparisonTest: Statistical comparison of two networks based on three invariance measures (2.2.2). R-CRAN. https://CRAN.R-project.org/package=NetworkComparisonTest

van Borkulo, C. D., van Bork, R., Boschloo, L., Kossakowski, J. J., Tio, P., Schoevers, R. A., Borsboom, D., & Waldorp, L. J. (2021). Comparing network structures on three aspects: A permutation test. Psychological Methods. doi:10.1037/met0000476.

van der Velden, P. G., Hyland, P., Contino, C., von Gaudecker, H. M., Muffels, R., & Das, M. (2021). Anxiety and depression symptoms, the recovery from symptoms, and loneliness before and after the COVID-19 outbreak among the general population: Findings from a Dutch population-based longitudinal study. PloS One, 16(1), e0245057. 10.1371/journal.pone.0245057

World Health Organization. (2019). International Classification of Diseases, Eleventh Revision (ICD-11). https://icd.who.int/

World Health Organization. (2022). Mental health and COVID-19: early evidence of the pandemic’s impact: scientific brief, 2 March 2022 CC BY-NC-SA 3.0 IGO). https://apps.who.int/iris/handle/10665/352189

Wysocki, A., van Bork, R., Cramer, A. O. J., & Rhemtulla, M. (2022). Cross-Lagged Network Models. 10.31234/osf.io/vjr8z

Zemestani, M., Beheshti, N., Rezaei, F., van der Heiden, C., & Kendall, P. C. (2021). Cognitive Behavior Therapy Targeting Intolerance of Uncertainty Versus Selective Serotonin Reuptake Inhibitor for Generalized Anxiety Disorder: A Randomized Clinical Trial. Behaviour Change, 38(4), 250–262. 10.1017/bec.2021.16

Zhang, Y., Hedley, F. E., Zhang, R. Y., & Jin, J. (2025). Toward quantitative cognitive-behavioral modeling of psychopathology: An active inference account of social anxiety disorder. J Psychopathol Clin Sci. 10.1037/abn0000972

