## Supplementary Material for "Transdiagnostic symptomatology amidst real-world environmental uncertainty: a cross-sectional and cross-lagged panel network analysis"

**Supplementary Table 1.** *Correlation matrix lower real-world uncertainty (t1).*

| | ANX | WOR | DEP | ANH | IUS1 | IUS2 | IUS3 | IUS4 | IUS5 | IUS6 | IUS7 | IUS8 | IUS9 | IUS10 | IUS11 | IUS12 |
|---|---|---|---|---|---|---|---|---|---|---|---|---|---|---|---|---|
| ANX | 1 | 0.819 | 0.595 | 0.446 | 0.432 | 0.314 | 0.302 | 0.347 | 0.333 | 0.342 | 0.267 | 0.415 | 0.411 | 0.379 | 0.393 | 0.362 |
| WOR | *** | 1 | 0.571 | 0.444 | 0.467 | 0.341 | 0.301 | 0.374 | 0.325 | 0.329 | 0.251 | 0.43 | 0.385 | 0.347 | 0.381 | 0.38 |
| DEP | *** | *** | 1 | 0.709 | 0.366 | 0.285 | 0.206 | 0.31 | 0.233 | 0.291 | 0.219 | 0.352 | 0.449 | 0.449 | 0.423 | 0.401 |
| ANH | *** | *** | *** | 1 | 0.368 | 0.241 | 0.154 | 0.298 | 0.201 | 0.28 | 0.155 | 0.36 | 0.379 | 0.372 | 0.423 | 0.4 |
| IUS1 | *** | *** | *** | *** | 1 | 0.494 | 0.429 | 0.612 | 0.445 | 0.532 | 0.359 | 0.528 | 0.516 | 0.517 | 0.456 | 0.534 |
| IUS2 | *** | *** | *** | *** | *** | 1 | 0.406 | 0.397 | 0.354 | 0.429 | 0.365 | 0.363 | 0.327 | 0.415 | 0.335 | 0.339 |
| IUS3 | *** | *** | *** | ** | *** | *** | 1 | 0.436 | 0.5 | 0.459 | 0.471 | 0.303 | 0.265 | 0.259 | 0.204 | 0.268 |
| IUS4 | *** | *** | *** | *** | *** | *** | *** | 1 | 0.459 | 0.541 | 0.382 | 0.467 | 0.48 | 0.481 | 0.445 | 0.467 |
| IUS5 | *** | *** | *** | *** | *** | *** | *** | *** | 1 | 0.496 | 0.529 | 0.431 | 0.373 | 0.385 | 0.36 | 0.409 |
| IUS6 | *** | *** | *** | *** | *** | *** | *** | *** | *** | 1 | 0.42 | 0.45 | 0.377 | 0.437 | 0.335 | 0.487 |
| IUS7 | *** | *** | *** | ** | *** | *** | *** | *** | *** | *** | 1 | 0.333 | 0.237 | 0.331 | 0.279 | 0.383 |
| IUS8 | *** | *** | *** | *** | *** | *** | *** | *** | *** | *** | *** | 1 | 0.682 | 0.619 | 0.636 | 0.641 |
| IUS9 | *** | *** | *** | *** | *** | *** | *** | *** | *** | *** | *** | *** | 1 | 0.786 | 0.738 | 0.632 |
| IUS10 | *** | *** | *** | *** | *** | *** | *** | *** | *** | *** | *** | *** | *** | 1 | 0.682 | 0.659 |
| IUS11 | *** | *** | *** | *** | *** | *** | *** | *** | *** | *** | *** | *** | *** | *** | 1 | 0.674 |
| IUS12 | *** | *** | *** | *** | *** | *** | *** | *** | *** | *** | *** | *** | *** | *** | *** | 1 |

*Note*: Pearson's correlations of network variables across samples. The correlation coefficient is on the upper diagonal and the strength of the *p*-value on the lower diagonal. ANX: anxiety/nervousness; WOR: worry; DEP: depression/hopelessness; ANH: anhedonia; IUS1-7: prospective IUS-subscale items; IUS8-12: inhibitory IUS-subscale items; IUS: Intolerance of Uncertainty Scale.
*** $p < .001$, ** $p < .01$.

**Supplementary Table 2.** *Correlation matrix higher real-world uncertainty (t2).*

| | ANX | WOR | DEP | ANH | IUS1 | IUS2 | IUS3 | IUS4 | IUS5 | IUS6 | IUS7 | IUS8 | IUS9 | IUS10 | IUS11 | IUS12 |
|---|---|---|---|---|---|---|---|---|---|---|---|---|---|---|---|---|
| ANX | 1 | 0.768 | 0.653 | 0.506 | 0.439 | 0.419 | 0.232 | 0.341 | 0.349 | 0.341 | 0.184 | 0.472 | 0.453 | 0.45 | 0.472 | 0.462 |
| WOR | *** | 1 | 0.64 | 0.557 | 0.457 | 0.449 | 0.257 | 0.406 | 0.29 | 0.362 | 0.121 | 0.45 | 0.437 | 0.47 | 0.442 | 0.473 |
| DEP | *** | *** | 1 | 0.727 | 0.401 | 0.332 | 0.216 | 0.363 | 0.297 | 0.339 | 0.15 | 0.418 | 0.429 | 0.444 | 0.439 | 0.428 |
| ANH | *** | *** | *** | 1 | 0.361 | 0.351 | 0.113 | 0.371 | 0.267 | 0.313 | 0.103 | 0.418 | 0.404 | 0.444 | 0.49 | 0.431 |
| IUS1 | *** | *** | *** | *** | 1 | 0.547 | 0.433 | 0.575 | 0.489 | 0.608 | 0.352 | 0.554 | 0.524 | 0.539 | 0.462 | 0.512 |
| IUS2 | *** | *** | *** | *** | *** | 1 | 0.41 | 0.405 | 0.427 | 0.45 | 0.316 | 0.418 | 0.361 | 0.422 | 0.358 | 0.409 |
| IUS3 | *** | *** | *** | n.s. | *** | *** | 1 | 0.446 | 0.396 | 0.427 | 0.476 | 0.279 | 0.252 | 0.288 | 0.178 | 0.338 |
| IUS4 | *** | *** | *** | *** | *** | *** | *** | 1 | 0.447 | 0.576 | 0.347 | 0.452 | 0.445 | 0.514 | 0.455 | 0.482 |
| IUS5 | *** | *** | *** | *** | *** | *** | *** | *** | 1 | 0.5 | 0.437 | 0.48 | 0.403 | 0.423 | 0.342 | 0.403 |
| IUS6 | *** | *** | *** | *** | *** | *** | *** | *** | *** | 1 | 0.42 | 0.493 | 0.428 | 0.489 | 0.376 | 0.502 |
| IUS7 | ** | * | ** | n.s. | *** | *** | *** | *** | *** | *** | 1 | 0.279 | 0.215 | 0.25 | 0.261 | 0.293 |
| IUS8 | *** | *** | *** | *** | *** | *** | *** | *** | *** | *** | *** | 1 | 0.624 | 0.622 | 0.612 | 0.619 |
| IUS9 | *** | *** | *** | *** | *** | *** | *** | *** | *** | *** | *** | *** | 1 | 0.773 | 0.724 | 0.679 |
| IUS10 | *** | *** | *** | *** | *** | *** | *** | *** | *** | *** | *** | *** | *** | 1 | 0.714 | 0.67 |
| IUS11 | *** | *** | *** | *** | *** | *** | *** | *** | *** | *** | *** | *** | *** | *** | 1 | 0.669 |
| IUS12 | *** | *** | *** | *** | *** | *** | *** | *** | *** | *** | *** | *** | *** | *** | *** | 1 |

*Note*: Pearson's correlations of network variables across samples. The correlation coefficient is on the upper diagonal and the strength of the *p*-value on the lower diagonal. ANX: anxiety/nervousness; WOR: worry; DEP: depression/hopelessness; ANH: anhedonia; IUS1-7: prospective IUS-subscale items; IUS8-12: inhibitory IUS-subscale items; IUS: Intolerance of Uncertainty Scale.
*** $p < .001$, ** $p < .01$, * $p < .05$, n.s. $p \geq .05$.

**Supplementary Table 3.** *Centralities for both networks.*

| | lower real-world uncertainty (*t*1) | | higher real-world uncertainty (*t*2) | |
|---|---|---|---|---|
| item | strength | bridge strength | strength | bridge strength |
| ANX | 0.532 | -0.554 | 0.253 | -0.019 |
| WOR | 0.649 | 0.309 | 1.044 | 0.758 |
| DEP | 0.285 | -0.812 | 0.147 | -1.606 |
| ANH | -1.474 | -0.129 | -0.547 | 0.655 |
| IUS1 | 1.185 | 2.114 | 1.249 | 0.884 |
| IUS2 | -2.006 | -0.506 | -1.554 | 0.096 |
| IUS3 | -0.65 | -1.429 | -0.255 | -0.866 |
| IUS4 | -0.345 | -0.127 | -0.507 | 0.296 |
| IUS5 | -0.217 | -0.382 | -1.032 | -0.231 |
| IUS6 | -0.097 | -0.186 | 0.356 | -0.47 |
| IUS7 | -1.423 | -1.133 | -2.171 | -1.707 |
| IUS8 | -0.059 | 1.284 | -0.182 | 2.006 |
| IUS9 | 1.532 | -0.563 | 0.66 | -1.302 |
| IUS10 | 1.003 | 0.737 | 1.384 | 0.463 |
| IUS11 | 0.392 | -0.366 | 0.946 | 0.276 |
| IUS12 | 0.693 | 1.742 | 0.209 | 0.769 |

*Note*: Standardised centrality indices of strength (i.e., sum of the absolute weights of every edge) and bridge strength (i.e., sum of absolute weights of edges across clusters) of cross-sectional networks at both timepoints (*t*1 and *t*2). ANX: anxiety/nervousness; WOR: worry; DEP: depression/hopelessness; ANH: anhedonia; IUS1-7: prospective IUS-subscale items; IUS8-12: inhibitory IUS-subscale items; IUS: Intolerance of Uncertainty Scale.

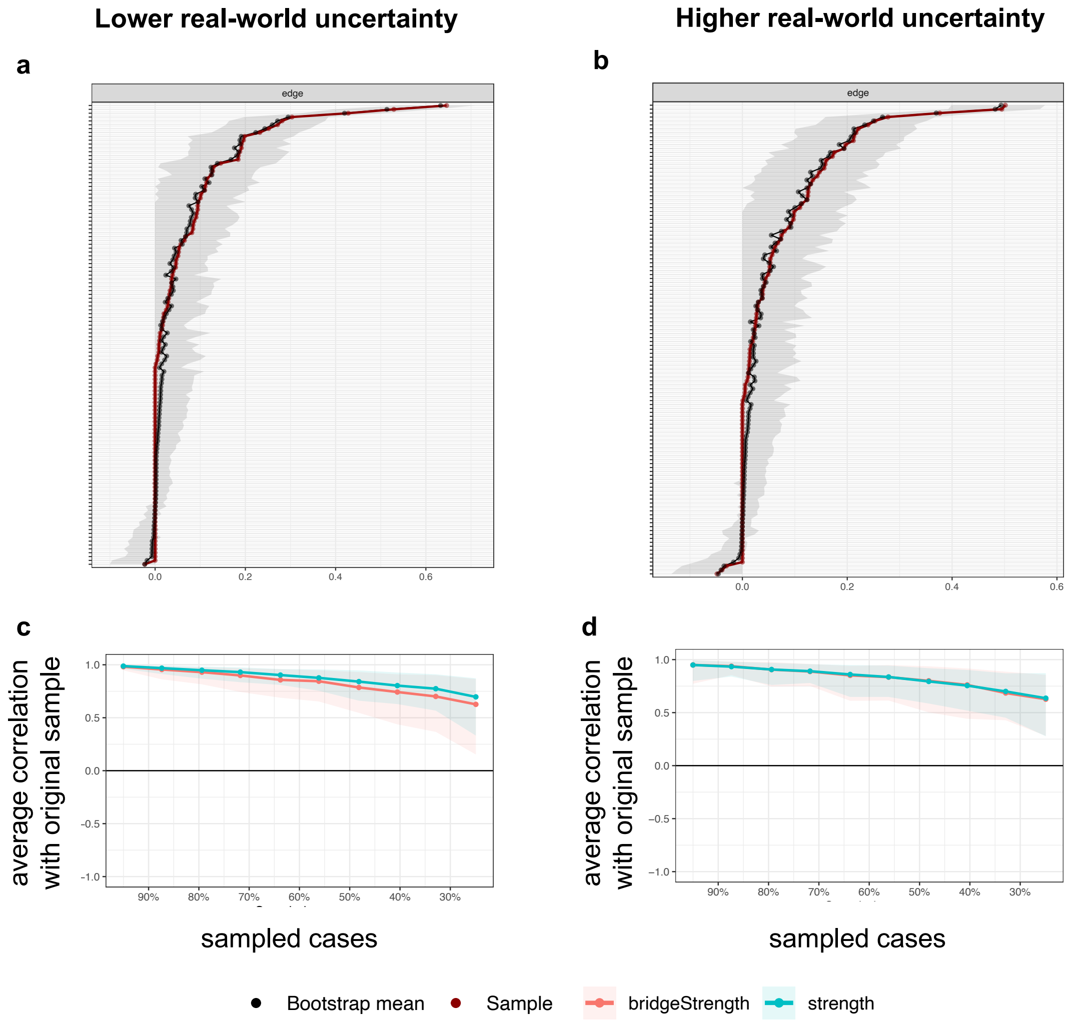


**Supplementary Figure 1. Bootstrapped stability for cross-sectional networks.**
**a** and **b** Accuracy of the edges weights established via non-parametric bootstrapped resampling for the two cross-sectional networks for **a** lower real-world uncertainty (*t*1) and **b** higher real-world uncertainty (*t*2). Red dots and lines: edge weights from the samples. Black dots and lines: edge weights generated based on 2000 random bootstrap samples. Gray shades: bootstrap-generated confidence intervals for each edge weight. **c** and **d** Correlation stability of centrality indices established via case-dropping bootstrapped resampling for **c** lower real-world uncertainty (*t*1) and **d** higher real-world uncertainty (*t*2). Solid lines link the means of correlations from subsets with increasing number of excluded cases. Colored areas indicate the range of correlations between 2.5th quantile and 97.5th quantile.

**Supplementary Table 4.** *Standardised regression matrix of CPLN model.*

| | higher real-world uncertainty (*t*2) | | | | | | | | | | | | | | | |
|---|---|---|---|---|---|---|---|---|---|---|---|---|---|---|---|---|
| lower real-world uncertainty (*t*1) | ANX | WOR | DEP | ANH | IUS1 | IUS2 | IUS3 | IUS4 | IUS5 | IUS6 | IUS7 | IUS8 | IUS9 | IUS10 | IUS11 | IUS12 |
| ANX | 0.289 | 0 | 0 | 0 | 0 | 0 | 0 | 0 | 0 | -0.058 | 0 | 0 | 0 | 0 | 0 | 0 |
| WOR | 0 | 0.343 | 0 | 0 | 0 | 0 | 0 | 0 | 0 | 0 | 0 | 0 | 0 | 0 | 0 | 0 |
| DEP | 0 | 0 | 0.413 | 0 | 0 | -0.179 | 0 | 0 | 0 | 0 | 0 | 0 | -0.08 | 0 | -0.066 | -0.083 |
| ANH | 0 | 0.082 | 0 | 0.296 | 0 | 0.164 | 0 | 0 | 0 | 0.086 | 0 | 0 | 0 | 0 | 0 | 0.062 |
| IUS1 | 0 | 0 | 0 | 0 | 0.699 | 0 | 0 | 0 | 0 | 0 | 0 | 0 | 0 | 0 | 0 | 0 |
| IUS2 | 0 | 0 | 0 | 0 | 0 | 0.724 | 0 | 0 | 0 | 0.051 | 0 | 0 | 0 | 0 | 0 | 0 |
| IUS3 | 0 | 0 | 0 | 0 | 0 | 0 | 0.763 | 0 | -0.117 | -0.066 | 0 | -0.142 | -0.101 | 0 | -0.069 | 0 |
| IUS4 | 0 | 0 | 0 | 0 | -0.081 | -0.096 | 0 | 0.62 | 0 | 0 | 0 | 0 | 0 | 0 | 0 | -0.062 |
| IUS5 | 0 | 0 | 0 | 0 | 0 | 0.078 | 0 | 0 | 0.837 | 0 | 0 | 0 | 0 | 0 | 0 | 0 |
| IUS6 | 0 | 0 | 0 | 0 | 0.073 | 0 | 0 | 0 | 0 | 0.821 | 0 | 0 | 0 | 0 | 0 | 0 |
| IUS7 | 0 | 0 | 0 | 0 | 0 | 0 | 0 | 0 | 0 | 0 | 0.566 | 0.11 | 0.068 | 0 | 0.079 | 0.063 |
| IUS8 | 0 | 0 | 0 | 0 | 0 | 0 | 0 | 0 | 0 | 0.061 | 0 | 0.643 | 0 | 0 | 0 | 0 |
| IUS9 | 0 | 0 | 0 | 0 | 0 | 0 | 0 | 0 | 0 | 0 | 0 | -0.13 | 0.746 | 0 | 0 | 0 |
| IUS10 | 0 | 0 | 0 | 0 | 0.117 | 0 | 0 | 0 | 0 | 0 | 0 | 0 | 0.114 | 0.794 | 0.125 | 0 |
| IUS11 | 0.221 | 0 | 0 | 0.171 | 0 | 0 | 0 | 0 | 0 | 0 | 0 | 0 | 0 | 0.096 | 0.782 | 0.072 |
| IUS12 | 0 | 0 | 0 | 0 | 0 | 0 | 0.096 | 0.108 | 0 | 0 | 0 | 0 | 0 | 0 | 0 | 0.813 |

*Note*: Edge weight results of the cross-lagged panel network from lower real-world uncertainty (rows) to higher real-world uncertainty (column). Diagonal items are autoregressive values. ANX: anxiety/nervousness; WOR: worry; DEP: depression/hopelessness; ANH: anhedonia; IUS1-7: prospective IUS-subscale items; IUS8-12: inhibitory IUS-subscale items; IUS: Intolerance of Uncertainty Scale.

**Supplementary Table 5.** *In-strength and out-strength in the CLPN network.*

| item | in-strength | out-strength |
|---|---|---|
| ANX | 0.221 | 0.058 |
| WOR | 0.082 | 0 |
| DEP | 0 | 0.409 |
| ANH | 0.171 | 0.395 |
| IUS1 | 0.271 | 0 |
| IUS2 | 0.517 | 0.051 |
| IUS3 | 0.096 | 0.495 |
| IUS4 | 0.108 | 0.239 |
| IUS5 | 0.117 | 0.078 |
| IUS6 | 0.322 | 0.073 |
| IUS7 | 0 | 0.32 |
| IUS8 | 0.382 | 0.061 |
| IUS9 | 0.364 | 0.13 |
| IUS10 | 0.096 | 0.356 |
| IUS11 | 0.339 | 0.559 |
| IUS12 | 0.343 | 0.204 |

*Note*: Indices of in-strength (predictability, i.e., sum of absolute weights of incoming edges) and out-strength (influence, i.e., sum of absolute weights of outgoing edges) of cross-lagged panel network (CLPN). ANX: anxiety/nervousness; WOR: worry; DEP: depression/hopelessness; ANH: anhedonia; IUS1-7: prospective IUS-subscale items; IUS8-12: inhibitory IUS-subscale items; IUS: Intolerance of Uncertainty Scale.
